# Architecture of the biofilm-associated archaic CupE pilus from *Pseudomonas aeruginosa*

**DOI:** 10.1101/2022.04.14.488289

**Authors:** Jan Böhning, Adrian Dobbelstein, Nina Sulkowski, Kira Eilers, Andriko von Kügelgen, Abul K. Tarafder, Vikram Alva, Alain Filloux, Tanmay A. M. Bharat

**Affiliations:** Sir William Dunn School of Pathology, University of Oxford, Oxford OX1 3RE, United Kingdom; Department of Protein Evolution, Max Planck Institute for Biology, Tübingen D-72076, Germany; Department of Life Sciences, MRC Centre for Molecular Bacteriology and Infection, Imperial College London, London SW7 2AZ, United Kingdom; Structural Studies Division, MRC Laboratory of Molecular Biology, Francis Crick Avenue, Cambridge CB2 0QH, United Kingdom

## Abstract

Chaperone-Usher Pathway (CUP) pili are major adhesins in Gram-negative bacteria, mediating bacterial adherence to biotic and abiotic surfaces. While classical CUP pili have been extensively characterized, little is known about so-called archaic CUP pili, which are phylogenetically widespread and promote biofilm formation by several human pathogens. In this study, we present the electron cryomicroscopy structure of the archaic CupE pilus from the opportunistic human pathogen *Pseudomonas aeruginosa*. We show that CupE pili consist of CupE1 subunits arranged in a zigzag architecture, with an N-terminal donor β-strand extending from each subunit into the next, where it is anchored by hydrophobic interactions, resulting in an overall flexible pilus arrangement. Imaging CupE pili on the surface of *P. aeruginosa* cells using electron cryotomography shows that CupE pili adopt variable curvatures in response to their environment, which may facilitate their role in promoting cohesion between bacterial cells. Finally, bioinformatic analysis shows the widespread abundance of *cupE* genes in isolates of *P. aeruginosa* and the co-occurrence of *cupE* with other *cup* clusters, suggesting interdependence of *cup* pili in regulating bacterial adherence within biofilms. Taken together, our study provides insights into the architecture of archaic CUP pili and their role in promoting cellular adhesion and biofilm formation in *P. aeruginosa*.

## Introduction

Adhesion of bacterial cells to abiotic and biotic surfaces is crucial for the colonization of new environments, including invasion of hosts during infections and biofilm formation (Klemm and Schembri, 2000, Soto and Hultgren, 1999, Hall-Stoodley et al., 2004, Berne et al., 2015, Melia et al., 2021). Bacterial adhesion is often mediated by proteinaceous, hair-like cell-surface structures known as pili or fimbriae (Nuccio and Baumler, 2007, Sauer et al., 2000). Pili are assembled by repeated interactions of monomeric protein subunits, leading to a filamentous structure that projects from the cell surface to anchor the cell to substrates (Fronzes et al., 2008).

<u>C</u>haperone-<u>U</u>sher <u>P</u>athway (CUP) pili are among the most widespread and well-characterized adhesins in Gram-negative bacteria (Waksman and Hultgren, 2009). CUP systems are typically encoded as single operons and consist of at least three components: a major pilin subunit, a periplasmic chaperone that stabilizes the pilin prior to assembly, and an outer membrane (OM) usher pore protein responsible for translocation and assembly of the pilin (Sauer et al., 2004, Barnhart et al., 2000, Dodson et al., 1993, Thanassi et al., 1998). Frequently, CUP operons encode further components, including adhesins that decorate the tip of the pilus distal to the OM (Lund et al., 1987), regulatory proteins, minor pilin subunits, and additional chaperones. CUP pilin subunits usually consist of an incomplete immunoglobulin-like (Ig-like) β-sandwich fold that lacks the final antiparallel β-strand, but contains an additional N-terminal β-strand extending away from the subunit core (Choudhury et al., 1999). In the assembled pilus, each pilin subunit provides its N-terminal β-strand to the following subunit to complete its Ig-like fold (Hospenthal et al., 2016) in a process termed donor strand complementation. Prior to assembly, the missing β-strand within the fold is donated by the chaperone protein (Choudhury et al., 1999). A tip adhesin subunit typically mediates the adhesive function of the pilus, often capping the pilus and mediating specific interactions with host receptors or other abiotic molecules (Jones et al., 1995, Krogfelt et al., 1990, Pakharukova et al., 2018). Additional pilin subunits encoded by many CUP operons typically fulfill a specialized structural role within the pilus (Jacob-Dubuisson et al., 1993, Lindberg et al., 1987, Kuehn et al., 1992) or have a role in terminating pilus assembly (Verger et al., 2006).

CUP pili are phylogenetically divided into three classes: classical, alternative, and archaic (Nuccio and Baumler, 2007). The best-characterized CUP pili belong to the classical type and include the Fim (Type 1 pilus) (Martinez et al., 2000, Jones et al., 1995) and Pap (P pilus) systems (Hospenthal et al., 2016, Lindberg et al., 1987) of *Escherichia coli*, which form stiff, tubular structures important for pathogenicity (Bouckaert et al., 2005, Martinez et al., 2000). However, structural information on non-classical CUP systems is scarce, and there have been no high-resolution studies of CUP pili in their cellular environment. Archaic CUP pili, the phylogenetically oldest class, are widespread in all proteobacteria, cyanobacteria, and even in some extremophilic phyla such as *Deinococcota* (Nuccio and Baumler, 2007). The best-characterized archaic CUP pili include the CupE system in *P. aeruginosa* (Giraud et al., 2011, Yen et al., 2002) and the Csu system in *Acinetobacter baumanii* (Tomaras et al., 2003, Pakharukova et al., 2018, Pakharukova et al., 2015). Both of these examples are from bacterial species belonging to the ESKAPE class of multidrug-resistant pathogens for which new antibiotics must urgently be developed (Pendleton et al., 2013), and targeting antimicrobials against pilins has been suggested as a potential therapeutic avenue (Ramezanalizadeh et al., 2020). Archaic CUP pili are crucial for the formation of biofilms (Tomaras et al., 2003, Giraud et al., 2011), which play an important role in persistent and chronic infections (Flemming et al., 2016, Davey and O’Toole, 2000).

In *P. aeruginosa*, the archaic CupE pilus is thought to have been acquired through horizontal gene transfer and evolved independently of other *cup* clusters in the genome (*cupA-D*), all of the latter belonging to the classical type of CUP systems (Giraud et al., 2011). The *cupE* gene cluster encodes one major pilin subunit (CupE1), two minor pilin subunits (CupE2, CupE3), a chaperone (CupE4), an usher (CupE5), and an adhesin protein (CupE6). The *cupE* operon is activated by a two-component system, PprA-PprB, and plays an important role in microcolony and macrocolony formation, as well as in maintaining the three-dimensional shape of the biofilm (Giraud et al., 2011, De Bentzmann et al., 2012). The expression of all CUP pili in *P. aeruginosa* is tightly regulated (Vallet et al., 2004), and the CupE pilus is expressed as part of an exopolysaccharide-independent adhesive signature together with the type I secretion system-dependent adhesin BapA, Type IVb pili and extracellular DNA (De Bentzmann et al., 2012). Hence, its function appears distinct from the other CUP systems in *P. aeruginosa* (CupA-D), which appear to be part of a different adhesive signature (Vallet et al., 2004, Vallet et al., 2001, Mikkelsen et al., 2009).

Comparatively less is known about the structure and architecture of archaic CUP pili, despite their presence in a wide range of species (Nuccio and Baumler, 2007). To bridge this critical gap in our knowledge, we purified CupE pili that were overproduced in *P. aeruginosa* cells through deletions of the CUP *mvaT* gene encoding a repressor, as well as PA2133, a phosphodiesterase encoded in the *cupA* operon. We used electron cryomicroscopy (cryo-EM) and cryotomography (cryo-ET) on natively assembled CupE pili *in vitro* and *in situ* to clarify their structural and architectural properties. We combined our structural and imaging experiments with bioinformatics, revealing important insights into the interdependence and role of CUP pili in biofilm formation of *P. aeruginosa*.

## Results

### Purification of natively assembled CupE pili

To study the role of CupE pili in *P. aeruginosa*, we engineered a strain with increased expression of CupE to facilitate isolation and structural analysis. Therefore, we utilized a previously described methodology of studying strains where other common cell surface filaments such as Type IV pili and flagella had been deleted (Vallet et al., 2001), which are otherwise abundant in pilus preparations. In addition, deletion of the gene encoding the MvaT repressor was performed, which had previously been shown to upregulate CUP pili in *P. aeruginosa* (Vallet et al., 2004). Thus, all experiments were performed using cells with a Δ*pilA* Δ*fliC* Δ*mvaT* background from hereon. During the bioinformatic analysis of *cup* operons in *P. aeruginosa*, we noted that the classical-type *cupA* operon also encodes a phosphodiesterase (PA2133, henceforth referred to as *cupA6*). As suggested by a previous study (Kulesekara et al., 2006), we reasoned that deletion of this gene could increase cellular cyclic di-guanylate (c-di-GMP) levels, a biofilm master regulator, and thus further increase the expression of CUP pili. Indeed, deletion of the *cupA6* phosphodiesterase resulted in increased expression of thin cell surface filaments (Figure S1A). This strain with increased surface filament expression was used to obtain preparations of purified filaments by shearing from the surface of cells scraped from plates (Methods). In the purified specimen, we observed long, curved filaments on cryo-EM grids (Figure 1A), with a different morphology to previously observed classical-type CUP filaments (Hospenthal et al., 2016), suggesting that these pili may correspond to archaic-type CupE pili rather than classical-type, tubular-shaped CupA-D pili. Indeed, mass spectrometric peptide fingerprinting confirmed that the sample contained CupE1 protein. These observations together suggest that combined deletion of genes encoding the MvaT repressor and the phosphodiesterase CupA6 caused increased expression of CupE filaments. To further verify that the observed filaments are CupE pili, we deleted *cupE1-E2* in the Δ*cupA6* background and found that the long, curved filaments with a ∼60 Å diameter were no longer present in fractions sheared from the cell surface (Figure S1B-C).

**Figure 1:**
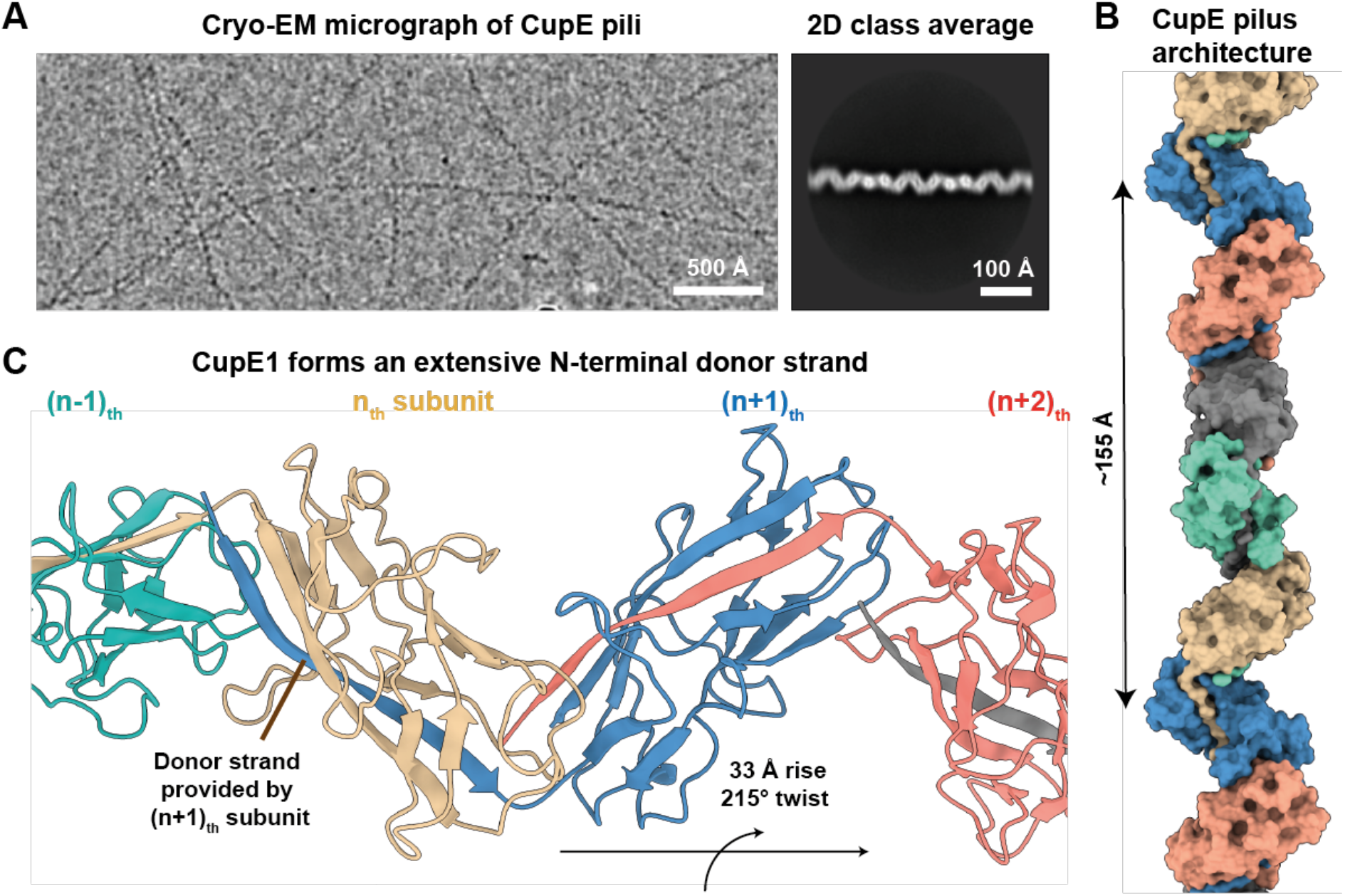
CupE1 subunits within the CupE pilus are arranged in a zigzag architecture. **(A)** Raw micrograph and 2D class average of CupE pili show zigzag-shaped filaments. **(B-C)** A 3.4 Å resolution cryo-EM structure of the CupE pilus reveals how CupE1 subunits are arranged in a zigzag pattern, with the donor strand of the (n+1)_th_ subunit enveloped by the incomplete Ig-like fold of each n_th_ subunit. Five CupE1 subunits form a longer repeat of ∼155 Å.

### Atomic structure of CupE pili using cryo-EM helical reconstruction

Having confirmed the presence of natively assembled CupE pili in our preparation, we proceeded with cryo-EM analysis. Cryo-EM micrographs and 2D classes of CupE showed a zigzag appearance of the pili (Figure 1A), with a diameter of approximately 60 Å. Interestingly, some pili spontaneously associated into large mesh-like bundles (Figure S1D). After deducing the symmetry of the pilus from two-dimensional class averages of single filaments (Figure 1A), we performed helical reconstruction and obtained a 3.5 Å resolution cryo-EM density, from which an atomic model of the CupE pilus could be built (Figures 1B-C and S2, Table S1, Movie S1). The atomic model reveals an arrangement of CupE1 subunits in a zigzag architecture (215° right-handed rotation per subunit, Figure 1B-C). In agreement with classical CUP proteins, the N-terminal β1-strand of each (n+1)_th_ subunit completes the Ig-like fold of the n_th_ subunit (Figure 2A-B). This donor strand interacts with the n_th_ subunit through both β-sheet hydrogen bonding as well as hydrophobic interactions, filling a hydrophobic groove in the subunit core (Figures 2C and S3A). The structure also shows two key cysteine residues (C41-C85) forming a disulfide bridge within a β-sheet from which the donor strand extends (Figure 2D). Bioinformatic analysis shows that these cysteines are highly conserved (Figure S4), suggesting they may play an important role in maintaining subunit stability.

**Figure 2:**
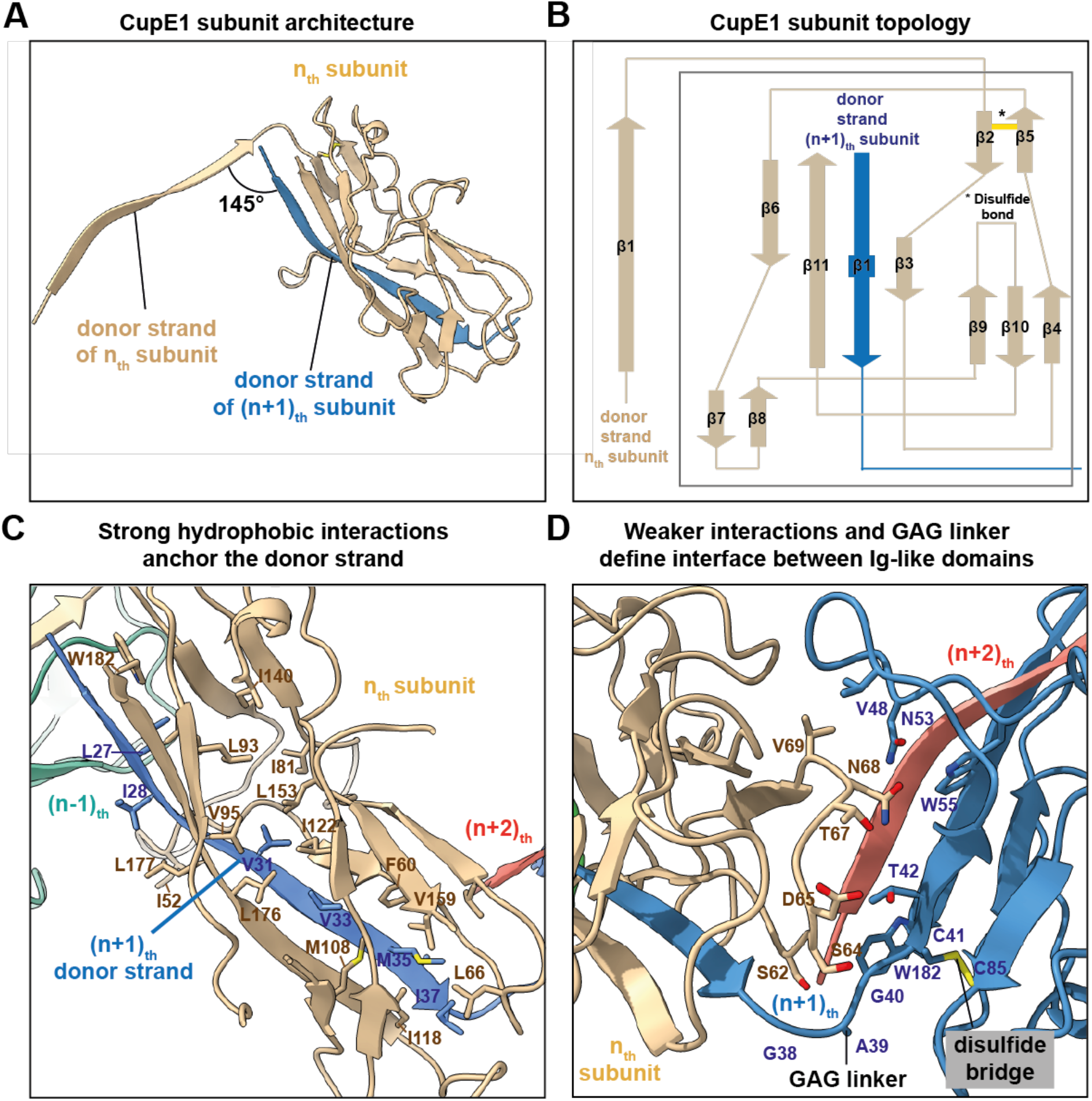
Architecture of the CupE1 subunit within the CupE pilus. **(A)** The donor strand from the (n+1)_th_ subunit completes a β-sheet in the n_th_ subunit by providing a 13-residue β-strand. **(B)** Subunit topology demonstrating β-sheet architecture of the CupE1 subunit. A yellow line denotes a disulfide bridge. **(C)** Extensive hydrophobic interactions anchor the donor strand of the (n+1)_th_ subunit into the Ig-like fold of the n_th_ subunit. **(D)** Compared to the extensive interactions of the donor strand with the complemented subunit, the globular subunit:subunit interface between Ig-like folds shows few strong interactions at the interface. Proximal to the donor strand, a GAG linker is positioning between adjoining subunits. A disulfide bridge is observed in each CupE1 subunit (marked).

The 13-residue donor strand of the (n+1)_th_ subunit represents the majority of the interaction with the n_th_ subunit, and the interface between the globular parts of Ig-like domains is relatively small (Figure 2D). Also, notably, the donor strand is connected to the core of each subunit through a Glycine-Alanine-Glycine (GAG) linker sequence (Figure 2D). Since GAG is a typical flexible linker motif (Robinson and Sauer, 1998), we hypothesized that flexibility around this hinge might be a key feature of the CupE pilus. Consistent with this hypothesis, curvature within the pilus was observed in a subset of two-dimensional class averages (Figure S3B).

A visual inspection of CupE pili along the helical axis (Figure S3B) showed a serine and threonine-rich loop (`TTTTSST`) extending outward from the helical axis of the CupE pilus, constituting the portion of the subunit that is most exposed to the environment (Figure S3C). Sequence alignment of archaic pilin subunits shows that this unusual serine and threonine-rich sequence is unique to *P. aeruginosa* (Figure S4). In the EM map, we noticed extra density near the loop that could not be attributed to the atomic model of the CupE1 protein subunit. Given that threonine and serine-rich sequences of bacterial adhesins are often post-translationally modified by O-glycosylation (Iwashkiw et al., 2013), we assigned this unexplained density as a potential glycan. Indeed, mutation of three threonine residues within the loop to Glycine-Alanine-Glycine, followed by cryo-EM structure determination led to a 4.1 Å resolution map with a smaller density in this region (Figure S3D-E, Table S1), confirmed by calculating difference maps with the wild-type pilus structure (Figure S3F), suggesting that this part of the CupE1 protein could be at least partially glycosylated.

### Imaging of CupE pili *in situ* on the surface of *P. aeruginosa* cells

CupE pili have been shown to promote the mature architecture and mushroom shape of *P. aeruginosa* biofilms (Giraud et al., 2011). To find out how CupE pili support cohesion between cells within the biofilm, we imaged CupE pili directly on the *P. aeruginosa* cell surface. To this end, we deposited cells from colonies of the Δ*cupA6* strain used above for obtaining pure CupE preparations onto grids for electron cryotomography (cryo-ET) imaging. In the resulting cellular electron cryotomograms (Figure 3), we observed CupE pili with the same characteristic zigzag architecture seen in *vitro* extending from the cell surface (Figures 3A-B and S5). To confirm that these were the same CupE pili, we extracted subtomograms along the length of the pili in tomograms and performed subtomogram averaging (Zivanov et al., 2022). Our subtomogram averaging structure of the cell surface filaments, produced from an unbiased cylindrical reference, recaptures the CupE pilus’ zigzag appearance (Figure S5), which was observed even without symmetrization, validating the identity of these filaments as CupE pili.

**Figure 3:**
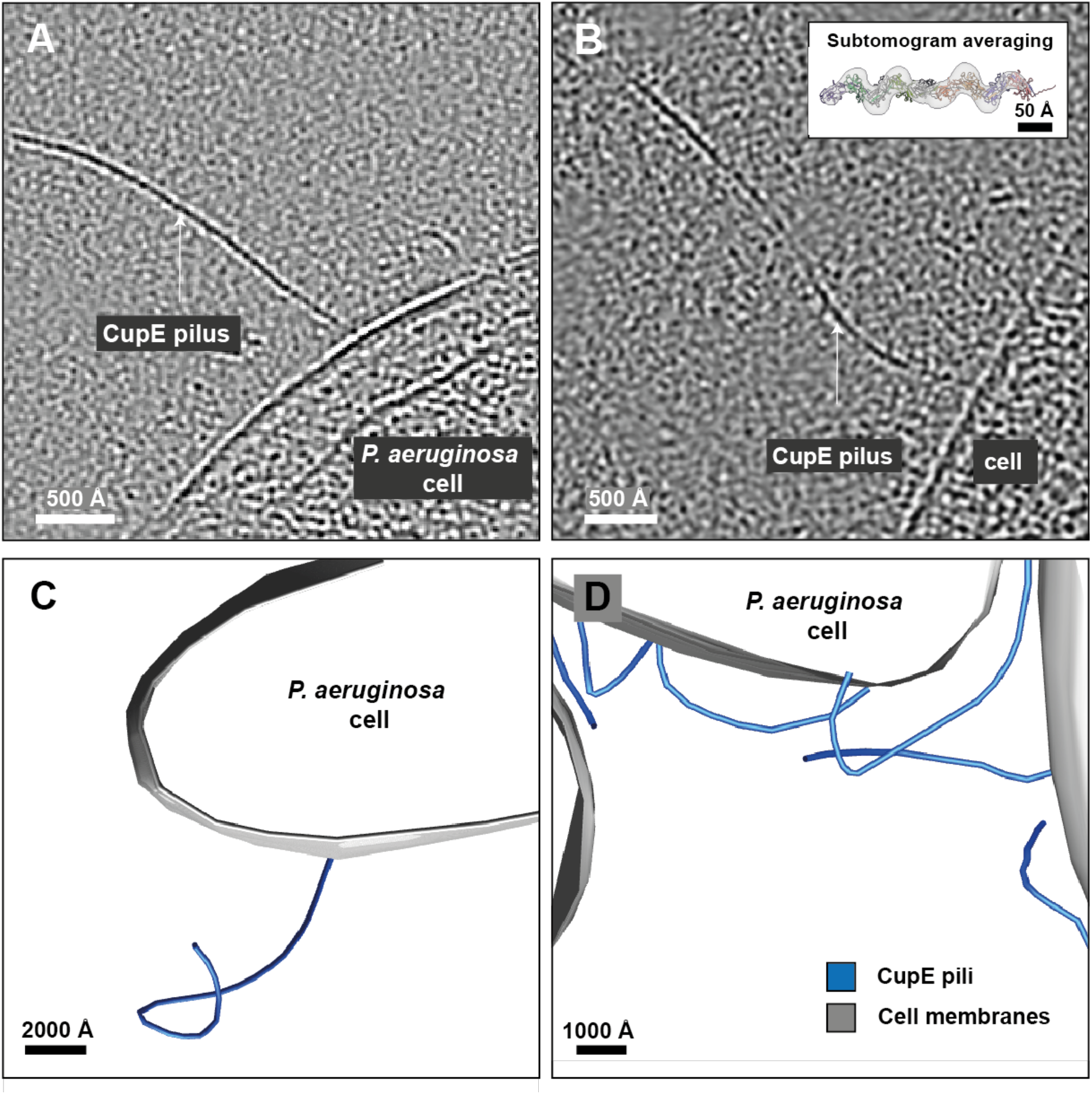
Cryo-ET imaging of CupE pili on *P. aeruginosa* cells. **(A-B)** Tomographic slice of cells expressing CupE pili. CupE pili adopt significant curvature on cells. The pilus can be seen going in and out of plane in (B). Inset: Atomic model of CupE fitted in the subtomogram averaging map produced from cellular cryo-ET data. **(C-D)** Segmentation of cell membranes and CupE pili illustrates how CupE pili adopt variable curvatures. See also Movie S2.

Interestingly, CupE pili attached to the cell adopted significantly variable curvatures *in situ* (Figures 3 and S5C-D, Movie S2). Varied curvature of CupE pili could also be observed within small clusters of cells, where pili were repeatedly found extending from and folding back onto the cells (Figure 3C-D). This observed ability of CupE pili to adopt variable curvature contrasts classical, tubular CUP pili, which are rigid assemblies (Hospenthal et al., 2016), and also contrasts observations made on the archaic Csu pilus, which is described to be inflexible as determined by EM and optical tweezer experiments (Pakharukova et al., 2021). Adopting varied curvature may be required for supporting efficient interactions of the pilus in the extracellular matrix of the biofilm, allowing the cells to embed themselves in a matrix rich in filamentous molecules. Here, flexibility of the pilus may help in promoting cell-cell adhesion within the biofilm.

### Structure- and sequence-based bioinformatic analysis of CupE pili

To place the structural and imaging data on CupE pili described above into the context of *P. aeruginosa* biofilm formation and CUP pili regulation, we performed computational analyses, facilitated by recent developments in protein structure prediction (Jumper et al., 2021) and methods for sequence analysis. While our cryo-EM structure is in agreement with previous studies that showed CupE1 to be the major CupE pilin subunit (Giraud et al., 2011), the *cupE* operon also encodes two additional predicted pilin subunits, CupE2 and CupE3, which exhibit high pairwise sequence similarity to CupE1, 39% and 30%, respectively (Figure S4). To model oligomers formed by the CupE2 and CupE3 subunits, we used AlphaFold-Multimer, which has been shown to predict the structures of monomeric and multimeric proteins with atomic-level accuracy (Evans et al., 2021). Our modelling suggests that CupE2 and CupE3 can form donor strand-exchanged filaments similar to CupE1 (Figure S6A). As a further verification of our modeling, we found that AlphaFold2 correctly predicted the CupE1 subunit arrangement consistent with our cryo-EM data (Figures S6 and S7), with minor subunit-subunit interface differences to our experimental structure. Finally, we also used AlphaFold2 to predict a structural model of the putative complex formed between CupE1 and the tip adhesin subunit CupE6. In the obtained model, as with the tip adhesin CsuE of *A. baumannii* (Pakharukova et al., 2018), CupE6 is arranged into two Ig-like folds. While the N-terminal domain is incomplete and caps the CupE1 filament by accepting a donor strand from CupE1, the distal end of the C-terminal fold contains a highly hydrophobic surface patch that likely plays a role in adhesion to target substrates (Figure S6C).

To investigate the conservation and co-occurrence of *cup* gene clusters in isolates of *P. aeruginosa* on a genomic level, we conducted exhaustive sequence searches of the *Pseudomonas* Genome Database (Winsor et al., 2016) and the NCBI RefSeq database. We searched for the occurrence of CupA-D and CupE systems with the usher as the query sequence (CupA3-CupD3 and CupE5), since the usher is the most conserved protein among the genes encoded by these clusters (Nuccio and Baumler, 2007). We detected complete *cupE* gene clusters, with preserved gene order, in 228 out of 233 strains of *P. aeruginosa*, showing the widespread occurrence of this archaic CUP cluster. Notably, *cupE* is missing in the strain PA7, which represents an extremely divergent isolate of *P. aeruginosa* (Weiser et al., 2019). Of the four classical *cup* clusters *in P. aeruginosa* (CupA-D), we identified the *cupB* cluster in all 233 strains, whereas *cupA* and *cupC* gene clusters were only missing in few strains (Table S5; Supplementary Data). Contrary to these, the *cupD* gene cluster, which had previously been found to be located on the pathogenicity island PAPI-1 (Mikkelsen et al., 2013, Mikkelsen et al., 2009), was only found in 11 strains. Interestingly, both the *cupA* and the *cupE* clusters are missing in most of the strains containing the *cupD* cluster. Given the high sequence similarity between the corresponding protein subunits of the CupA and CupD systems (>65% pairwise sequence identity), we speculate they may fulfill highly similar functions and are, therefore, often mutually exclusive. Taken together, three different classical *cup* gene clusters and one archaic gene cluster were found to be widespread in *P. aeruginosa* strains.

The success of our strategy to upregulate CupE pili by deleting a phosphodiesterase-encoding gene in the *cupA* operon (*cupA6)* suggests that CUP pili expression may be interdependent on other CUP genes, implying potential co-regulation. This agrees with the findings of our bioinformatics analysis, as CupA and CupE co-occur in almost all strains of *P. aeruginosa*. Notably, the phosphodiesterase CupA6 is encoded after the chaperone CupA5 with a sequence overlap of four nucleotides in every strain containing CupA (Figure 4A), indicating likely translation coupling between them (McCarthy and Gualerzi, 1990).

**Figure 4:**
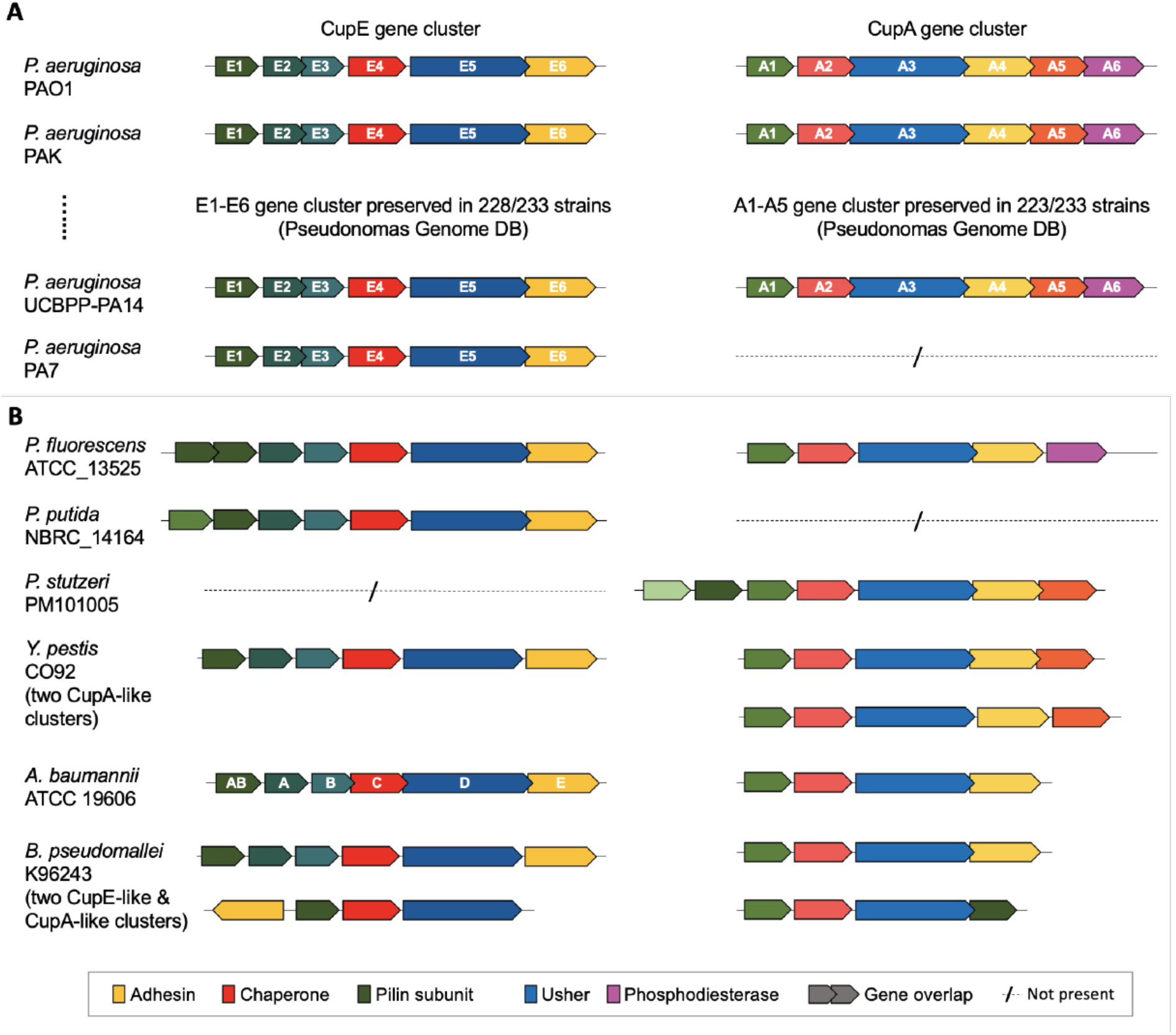
Co-occurrence of *cupA* and *cupE* gene clusters. **(A)** Bioinformatic analysis reveals co-occurrence of the *cupA* and *cupE* gene clusters with preserved gene order in most *P. aeruginosa* strains. Overlapping sequences of different CUP genes indicates possible translational coupling; of the usher gene *cupE5* to the adhesive tip subunit gene *cupE6*, of the two minor pilin subunits *cupE2* and *cupE3*, and of all genes *cupA2*-*cupA6*. A phosphodiesterase (CupA6) is encoded after *cupA5* in every strain in which *cupA* occurs. **(B)** Outside *P. aeruginosa*, examples of co-occurrence can be found in pathogenic proteobacteria such as *P. fluorescens, Y. pestis, A. baumannii*, and *B. pseudomallei*. Gene order in CUP clusters outside *P. aeruginosa* is not preserved; e.g. in *P. fluorescens* ATCC_13525, an additional pilin is encoded, and in *B. pseudomallei* K96243, the *cupA6-*like gene is encoded on the opposite strand. Accession data and gene loci are provided in Table S5 and as Supplementary Data.

Outside *P. aeruginosa, cup* gene clusters are more divergent and do not share a preserved gene order, exhibiting missing, additional, or swapped genes. In other bacteria of the genus *Pseudomonas, cupE*-like gene clusters are more widespread than *cupA*-*cupD* and co-occurrence of *cupA* and *cupE* is rare (Figure 4); for instance, while *P. putida* NBRC_14164 lacks *cupA, P. fluorescens* ATCC_13525 contains both *cupA* and *cupE*. Co-occurrence of *cupE*-like and *cupA*-like gene clusters were also identified in some species of other gammaproteobacteria (e.g., *Yersinia pestis, A. baumannii*) as well as betaproteobacteria (e.g., *Burkholderia pseudomallei*) (Figure 4B).

## Discussion

In this study, we present the structure of an archaic CUP pilus, demonstrating a zigzag architecture with extensive interactions between the donor strand and the complemented subunit. The subunit interface between the globular part of the Ig-like domains has significantly fewer inter-subunit contacts, suggesting that flexibility around this interface may be a feature of the archaic CUP pilus. Indeed, imaging of CupE pili *in situ* shows they can adopt significantly varied curvatures, which may aid in interactions with the biofilm matrix. This shows that the CupE pilus is not a stiff, tubular assembly, but more akin to a rope, which may ‘wrap’ around other objects while maintaining adhesion to its target. The observed lateral flexibility of the CupE pilus contrasts with the Csu pilus from *A. baumanii*, which, in a recent study employing overexpression of the corresponding *csu* operon in *E. coli* (Pakharukova et al., 2021), has been observed to form stiff filaments, although the pilus was found to be able to extend to almost twice its length in optical tweezer experiments. These results and those from our study indicate the likely possibility that the interface between the Ig-like domains of the archaic CUP pilus enables structural flexibility, and may enable either bending (this study) or stretching (Pakharukova et al., 2021).

AlphaFold structure predictions of minor CupE subunits CupE2 and CupE3 suggest that they too form filaments through donor strand exchange. However, it is unclear whether their purpose is to functionalize pili or fulfil an undetermined function in pilus assembly. For example, in the Pap system, the PapH subunit terminates pilus assembly (Båga et al., 1987, Verger et al., 2006); whether an analogous protein exists in archaic CUP pili remains to be determined. Structural prediction of the CupE6 adhesin tip subunit also suggests that, in agreement with previous analyses on the Csu system (Pakharukova et al., 2018), CupE6 contains the same subunit fold and hydrophobic surface at the tip that is thought to interact with other hydrophobic components, thus supporting adhesion. The chemical nature of the substrate of the CupE6 pilus tips in biofilms remains enigmatic, and it is unclear how the highly hydrophobic pilus tip would be stabilized during pilus assembly, presenting an exciting direction for future inquiries.

CupE and Csu, the two archetypal archaic CUP pili systems, are both involved in promoting biofilm formation. Our study shows that the flexibility of such pili may be a key feature helping them adapt to the complex three-dimensional biofilm environment, allowing them to promote biofilm formation efficiently. While we study CupE pili on cells, an intrinsic limitation of our system is that single cells expressing CupE do not fully recapture the molecular sociology and crowding conditions within intact biofilms. Interaction partners of the pilus are hence unknown and warrant the focus of future imaging efforts. Since we have observed isolated CupE pili forming regular mesh-like arrays in cryo-EM images, this might indicate that one such interaction partner may be other CupE pili (Figure S1D), hinting that lateral interactions of pili may occur in the crowded conditions of the biofilm matrix, similar to other biofilm matrix fiber systems (Tarafder et al., 2020, Boehning et al., 2022). Further studies on native cellular systems – i.e., biofilms - will be required to determine the exact mode of interaction of CupE pili with other extracellular matrix components and cells.

Moreover, our study finds that deletion of the gene encoding the phosphodiesterase CupA6, which is encoded in the *cupA* operon, results in increased expression of the *cupE* operon. A likely explanation for this is that the *cupA* operon negatively regulates cyclic di-GMP levels through CupA6, and that deletion of CupA6 causes higher di-GMP levels, subsequently causing *cupE* expression. Indeed, we find that *cupA* and *cupE* gene clusters mostly co-occur in *P. aeruginosa* isolates, suggesting both fulfill a distinct function during biofilm development, and that their expression may be co-regulated and interdependent. Numerous CUP systems and other adhesins have been identified in *P. aeruginosa*, and many have been found to be of general importance for biofilm formation. This prompts the overarching question: Why does *P. aeruginosa* have, in the same manner as a Swiss army knife, an arsenal of different adhesins? Are some adhesin systems co-operative or expressed in specialized conditions - and if so, when? Answering these questions in future studies will greatly enhance our understanding of adhesion, biofilm formation, and pathogenicity of *P. aeruginosa* specifically and Gram-negative bacteria in general.

## Materials and Methods

### Construction of *P. aeruginosa* mutants

*P. aeruginosa* deletion mutants were created as described previously utilising the suicide plasmid pKNG101 (Muhl and Filloux, 2014). Briefly, to engineer gene deletions in the PAO1 strain, 500 bp DNA fragments of the 5’ (upstream) and 3’ (downstream) ends of the gene of interest were obtained by PCR using PAO1 chromosomal DNA as a template. The upstream fragment was amplified with the oligonucleotides P1 and P2 while the downstream fragment was amplified using P3 and P4 (Table S2). A third PCR step using P1 and P4 resulted in a DNA fragment with the flanking region of the gene of interest. The gene fragment was then cloned into pCR-BluntII-TOPO (Invitrogen), the sequence confirmed and sub-cloned into the pKNG101 suicide vector (Table S3). The pKNG-derivatives were maintained in *E. coli* strain CC118λpir (for strain descriptions, see Table S4) and conjugated into PAO1 using *E. coli* 1047 harbouring the conjugative plasmid pRK2013. pKNG101 was conjugated into *P. aeruginosa* as described in (Muhl and Filloux, 2014). After homologous recombination, colonies were streaked onto agar containing 20% (w/v) sucrose and grown at room temperature for 48 hours to select for colonies that have lost the plasmid backbone. Gene deletions were verified by PCR using external primers P5 and P6 (Table S2). Chromosomal substitution of TTTTSST to AGATSST in CupE1 was achieved by reintroduction of the previously deleted *cupE1-2* fragment into PAO1 Δ*pilA* Δ*fliC* Δ*mvaT* Δ*cupA6* Δ*cupE1-2*. In brief, the *cupE1-2* DNA fragment was amplified using primers called Δ*cupE1-2* P1 and P4 (Table S2), subcloned into pCR-BluntII-TOPO (Invitrogen) and subjected to site directed mutagenesis using CupE1-AGATSST Fw and Rev primers (Table S2). After the sequence was confirmed, the fragment was cloned into pKNG101 and conjugation was conducted as described above. Mutation was verified by PCR using external primers Δ*cupE1-2* P5 and P6 and clones with *cupE1-2* fragment reintroduced were sequenced.

### Isolation of CupE pili

An overnight culture of *P. aeruginosa* PAO1 Δ*mvAT* Δ*cupA6* Δ*pilA* Δ*fliC*, grown in lysogeny broth (LB) medium at 37°C with agitation at 180 revolutions per minute (rpm), was used to plate lawns on agar plates, and incubated overnight at 37 °C. Bacterial lawns were scraped and resuspended in 1x phosphate-buffered saline (PBS). The resulting suspension was vortexed for 90 seconds to promote dissociation of pili from the cell surface. Cells were then centrifuged at 4,500 relative centrifugal force (rcf) for 20 minutes, and the supernatant centrifuged again 3-4 times at 16,000 rcf to remove remaining cells and cellular debris. 500 mM NaCl and 3% (w/v) PEG-6000 were added to the supernatant, and pili were precipitated on ice for 1 hour. Precipitated pili were collected via centrifugation for 30 minutes at 16,000 rcf. For cryo-EM, precipitated pellets were combined and precipitated again in the same manner and resuspended in 1x PBS to produce the final product.

### Negative stain electron microscopy

2.5 µl of sample was applied to a glow-discharged carbon support grid (TAAB), blotted, washed three times with water, and stained using three 20 µl drops of 2% (w/v) uranyl acetate and allowed to air-dry. Negatively stained grids were imaged on a Tecnai T12 microscope.

### Cryo-EM and cryo-ET sample preparation

For cryo-EM grid preparation of purified CupE pili, 2.5 µl of the sample was applied to a freshly glow-discharged Quantifoil R 2/2 Cu/Rh 200 mesh grid and plunge-frozen into liquid ethane using a Vitrobot Mark IV (ThermoFisher) at 100% humidity at an ambient temperature of 10 °C. For tomography sample preparation of PAO1 Δ*pilA* Δ*fliC* Δ*mvaT* Δ*cupA6*, a bacterial lawn from an overnight LB agar plate incubated at 37 °C without antibiotics was resuspended in PBS, and 10 nm Protein-A-gold beads (CMC Utrecht) were added as fiducials prior to plunge-freezing.

### Cryo-EM and cryo-ET data collection

Cryo-EM data was collected in a Titan Krios G3 microscope (ThermoFisher) operating at an acceleration voltage of 300 kV, fitted with a Quantum energy filter (slit width 20 eV) and a K3 direct electron detector (Gatan). Images were collected in super-resolution counting mode using a physical pixel size of 1.092 Å/pixel for helical reconstruction of CupE pili and 3.489 Å/pixel for cellular tomography data. For helical reconstruction of CupE, movies were collected as 40 frames, with a total dose of 45-46 electrons/Å^2^, using a range of defoci between -1 and -2.5 µm. For the wild-type CupE pilus dataset, 11,584 movies were collected; for the 111-113_AGA_ dataset, 4,665 movies were collected. Cryo-ET tilt series of PAO1 Δ*pilA* Δ*fliC* Δ*mvAT* Δ*cupA6* cells were collected using a dose-symmetric tilt scheme as implemented in SerialEM (Mastronarde, 2005), with a total dose of 121 electrons/Å^2^ per tilt series and defoci of -8 to -10 µm, and with ±60º tilts of the specimen stage at 1º tilt increments.

### Cryo-EM processing

Helical reconstruction of CupE pili was performed in RELION 3.1 (He and Scheres, 2017, Zivanov et al., 2020, Scheres, 2012). Movies were motion-corrected and Fourier-cropped using the RELION 3.1 implementation of MotionCor2 (Zheng et al., 2017), and CTF parameters were estimated using CTFFIND4 (Rohou and Grigorieff, 2015). Initial helical symmetry of CupE pili was estimated through indexing of layer lines and counting the number of visible subunits along the pilus. Three-dimensional classification was used to identify a subset of particles that supported refinement to 3.5 Å resolution. For final refinement, CTF multiplication was used for the final polished set of particles (Zivanov et al., 2018, Zivanov et al., 2020, Zivanov et al., 2019). Symmetry searches were used during reconstruction, resulting in a final rise of 33.12 Å and a right-handed twist per subunit of 214.56°. Resolution was estimated using the gold-standard Fourier Shell Correlation (FSC) method as implemented in RELION 3.1. Local resolution measurements were also performed using RELION 3.1.

### Model building and refinement

Manual model building of the CupE1 subunit was performed in Coot (Emsley et al., 2010) as follows. A homology model based on the structure of *A. baumanii* CsuA/B (RCSB 6FQA) was calculated using MODELLER (Webb and Sali, 2016) and this homology model was fit into the cryo-EM density as a rigid body. Residues of the homology model that were inconsistent with the density, including the N-terminal donor strand, were deleted and manually rebuilt. The initially built model was subjected to real-space refinement against the cryo-EM map within the Phenix package (Adams et al., 2010, Afonine et al., 2018). Five subunits of CupE1 were built and used for final refinement. Non-crystallographic symmetry between individual CupE1 subunits was applied for all refinement runs. Model validation including map-vs-model resolution estimation was performed in Phenix.

### Tomogram reconstruction

Tilt series alignment via tracking of gold fiducials was performed using the etomo package as implemented in IMOD (Kremer et al., 1996). Tomograms were reconstructed with WBP in IMOD or SIRT in Tomo3D (Agulleiro and Fernandez, 2015, Fernandez et al., 2018). Deconvolution of tomograms using the tom_deconv.m script (Tegunov and Cramer, 2019) was performed for visualisation purposes.

### Subtomogram averaging

Subtomogram averaging of pili on cells was performed in RELION 4 (Zivanov et al., 2022), employing helical reconstruction (He and Scheres, 2017). A cylindrical reference was used to avoid bias. Helical symmetry was applied to enhance the signal during particle alignment. The map presented in Figures 3 and S5 is unsymmetrized.

### Data visualisation and quantification

Cryo-EM images were visualized in IMOD. Fiji (Schindelin et al., 2012) was used for bandpass and Gaussian filtering, followed by automatic contrast adjustment. Atomic structures and tomographic data were displayed in ChimeraX (Goddard et al., 2018). Segmentation of tomograms was performed manually in IMOD. Quantification of cell surface filaments in the Δ*cupA6* mutant was performed through manual annotation in 30 randomly acquired negative stain EM images targeted on cells located at low magnification. Atomic models are shown in perspective view, except for Figure S3C, which is shown in orthographic view. Hydrophobic surfaces were calculated in ChimeraX using the in-built *mlp* function. Difference maps were calculated using EMDA (Warshamanage et al., 2022) with maps lowpass-filtered to the same resolution (4.2 Å).

### Bioinformatic analysis

Sequence data was downloaded from the Pseudomonas Genome Database v. 20.2 (Winsor et al., 2016) and filtered to exclude incomplete genomes. Searches were performed against every single strain using PSI-BLAST (Camacho et al., 2009), with the proteins CupA3 (PA2130 NCBI locus tag), CupB3 (PA4084), CupC3 (PA0994), and CupE5 (PA4652) from the reference strain *P. aeruginosa* PAO1 as queries. Because the CupD system is missing in the strain PAO1, the CupD3 usher (PA14_59735) from strain UCBPP-PA14 was used as the query. To get a unique assignment of genes to CUP proteins, output data was further filtered with custom scripts and probable sequencing errors were corrected.

Structure predictions were conducted using AlphaFold-Multimer version 2.1.1 (Jumper et al., 2021, Evans et al., 2021), with sequences from the reference genome PAO1 as queries. Filaments were predicted without the signal peptide, which was predicted using SignalP-6.0 (Teufel et al., 2022). The multiple sequence alignments (MSAs) used for the structure inference were built with the standard AlphaFold pipeline and the “reduced_dbs” preset. Template modelling was enabled and structures were inferred with one MSA recycling iteration and all five different model parameter sets. After prediction, models were ranked by the pTM score and only the highest-ranking model was selected. PAE-value-plots for each structure are shown in Figure S7. All predictions were performed using the high-performance computer “Raven”, operated by the Max-Planck Computing & Data facility in Garching, Munich, Germany. The multiple sequence alignment shown in Figure S4 was obtained by first calculating an initial alignment using PROMALS3D (Pei et al., 2008) in default settings and by subsequently curating it manually.

## Acknowledgments

T.A.M.B. is a recipient of a Sir Henry Dale Fellowship, jointly funded by the Wellcome Trust and the Royal Society (202231/Z/16/Z). T.A.M.B. would like to thank the Vallee Research Foundation, the Leverhulme Trust and the Lister Institute of Preventative Medicine for support. J.B. is supported by a Medical Research Council graduate studentship (grant numbers MR/K501256/1 and MR/N013468/1). The authors would like to thank Dr. Thomas Clamens for help with strain generation.

## Author Contributions

J.B., N.S., V.A., A.F. and T.A.M.B. designed research. J.B., A.D., N.S., K.E., A.K.T., A.v.K., V.A. and T.A.M.B. performed research. J.B., A.D., N.S., A.K.T., A.v.K., V.A. and T.A.M.B. analysed data. J.B., A.D., K.E., V.A., A.F. and T.A.M.B wrote the manuscript with support from all the authors.

## Figures

**Figure S1:**
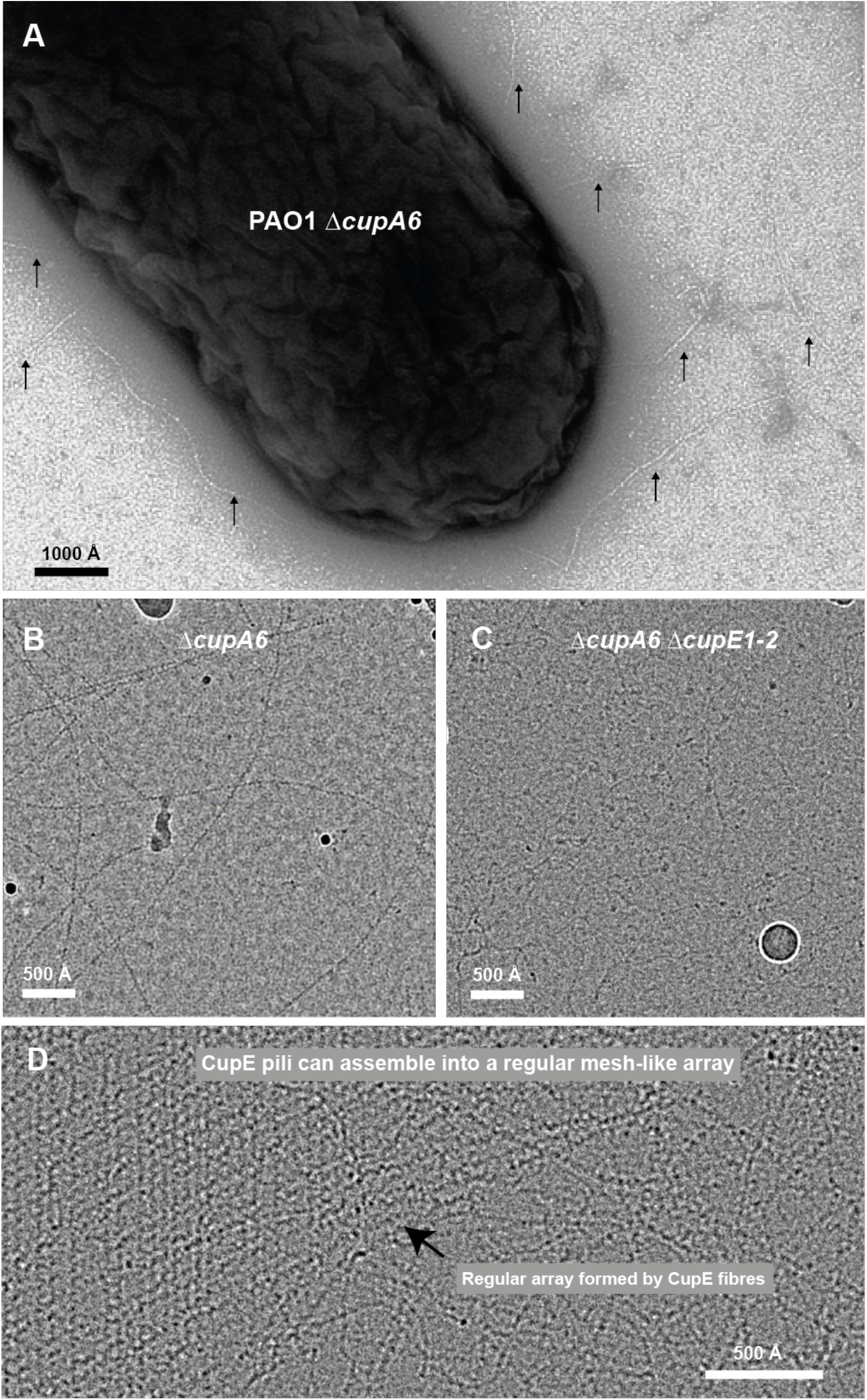
Deletion of *cupA6* causes increased expression of CupE pili. **(A)** Negative stain image of a *P. aeruginosa* PAO1 strain with *cupA6* deletion. Deletion of *cupA6* causes upregulation of cell-surface fibers compared to a control strain without the deletion (188 fibers in the Δ*cupA6* strain versus 60 fibers in a control strain; n=30 micrographs; see Methods). Fibers are highlighted by arrows. **(B-C)** Cell surface filaments were sheared, precipitated and subjected to cryo-EM, showing (B) the Δ*cupA6* strain as shown in (A), and (C) a Δ*cupA6* Δ*cupE1-2* strain, demonstrating that the cell surface pili with a dotted zigzag pattern are CupE pili. Small fiber contamination is enriched in (C). This contaminant is possibly DNA due to its size and low persistence length, which precipitates under similar conditions (Paithankar and Prasad, 1991). **(D)** In the cryo-EM dataset of purified CupE pili, instances of CupE pili forming a crisscross mesh-like array were observed.

**Figure S2:**
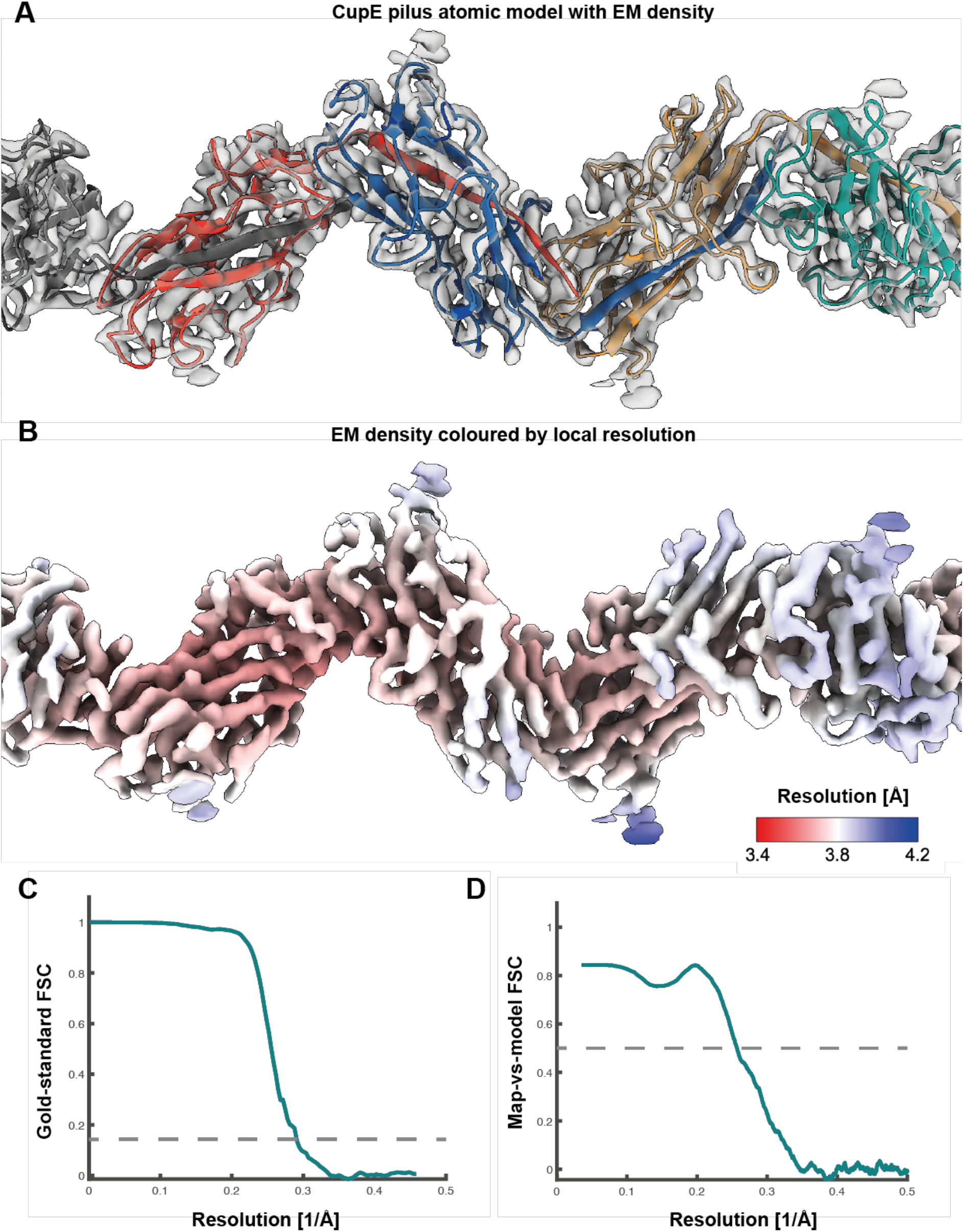
Cryo-EM of the CupE pilus. **(A)** Atomic model of the CupE pilus, consisting of CupE1 subunits, in the transparent cryo-EM density at 15 σ away from the mean. **(B)** The same density shown in (A) coloured according to local resolution. **(C)** Gold-standard FSC curve. **(D)** Map-vs-model FSC curve.

**Figure S3:**
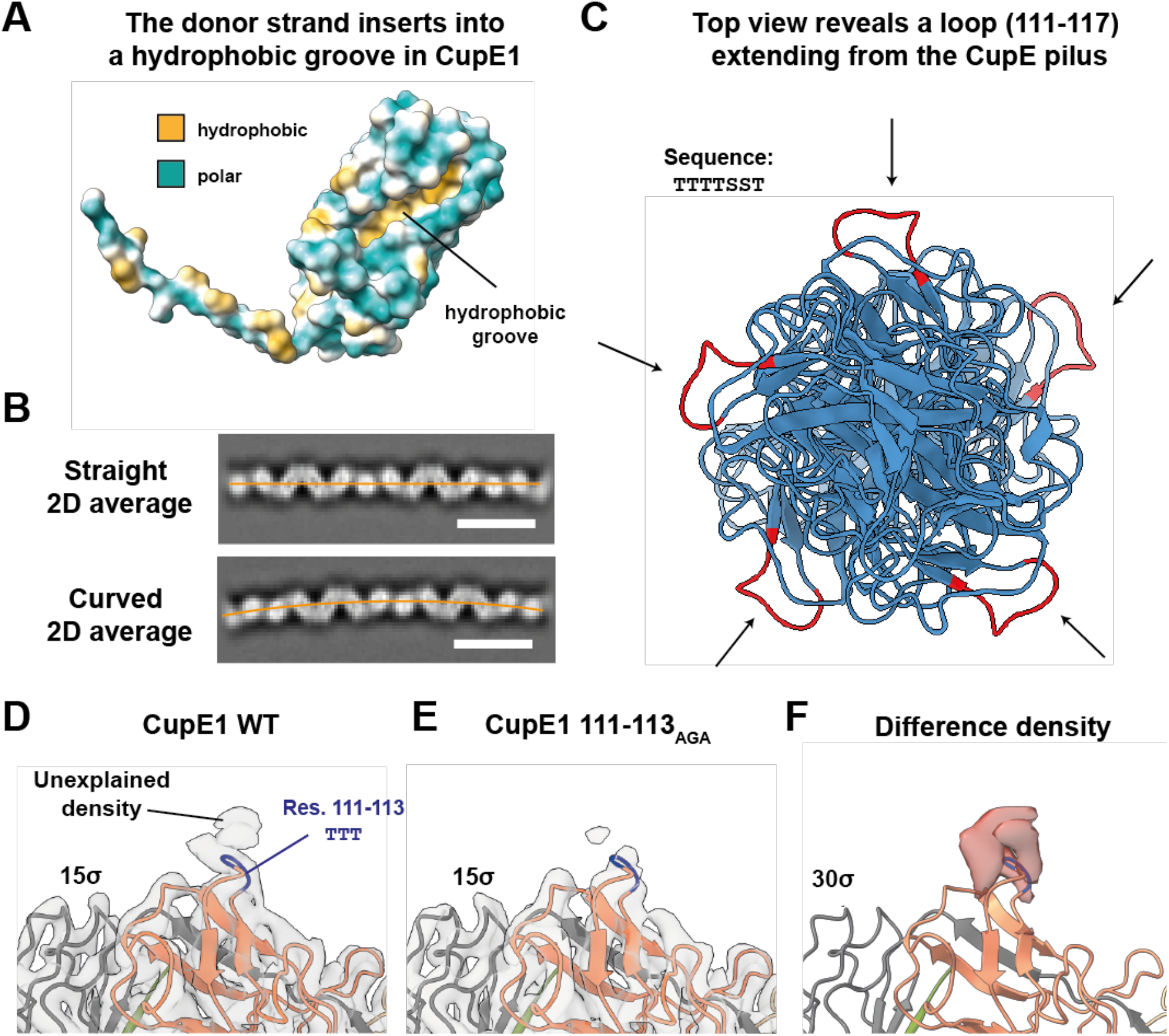
Structural features of the CupE pilus. **(A)** Hydrophobic surface depiction of an uncomplemented CupE1 subunit reveals that the donor strand inserts into a hydrophobic groove. **(B)** Straight and curved 2D class averages of CupE pili. Orange lines indicate the center of the filament to facilitate the visualization of curvature. Scale bars are 100 Å. **(C)** Top view of a five-subunit ribbon model of the CupE pilus reveals that a serine-threonine-rich loop (marked red, sequence `TTTTSST`) extends from the pilus, exposed to the environment. **(D-F)** Mutation of the first three residues of the loop shown in (C) to AGA (111-113_AGA_) followed by structural determination at 4.1 Å resolution via cryo-EM shows reduced density near the loop, suggesting this density could arise from post-translational modifications. Density is shown at 15 σ contour level in (D) and (E), difference density is shown at 30 σ in (F).

**Figure S4:**
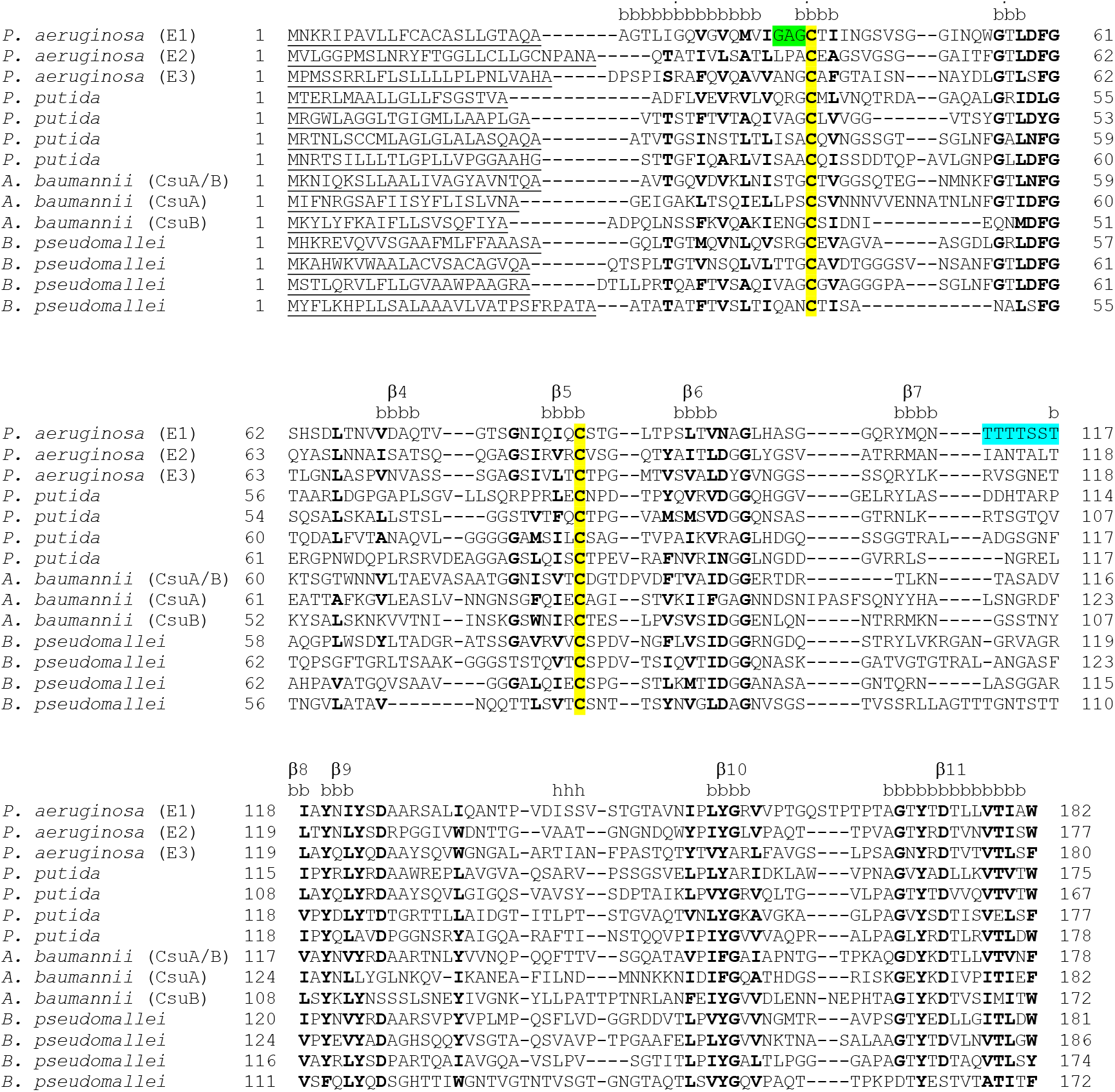
Multiple sequence alignment of major and minor pilin subunits of archaic CUP pili. Conserved residues are shown in boldface. Secondary structure (b=β-strand, h=*α*-helix) is annotated based on our cryo-EM structure of the CupE1 filament. The conserved cysteine residues (C41 and C85) that form a disulfide bond in CupE1 filaments are highlighted in yellow, the ‘GAG’ linker in green, and the serine-threonine-rich loop in cyan. The signal peptide in each sequence, as predicted using SignalP 6.0 (Teufel et al., 2022), is underlined. Accession details for the shown protein sequences are provided in the Table S5.

**Figure S5:**
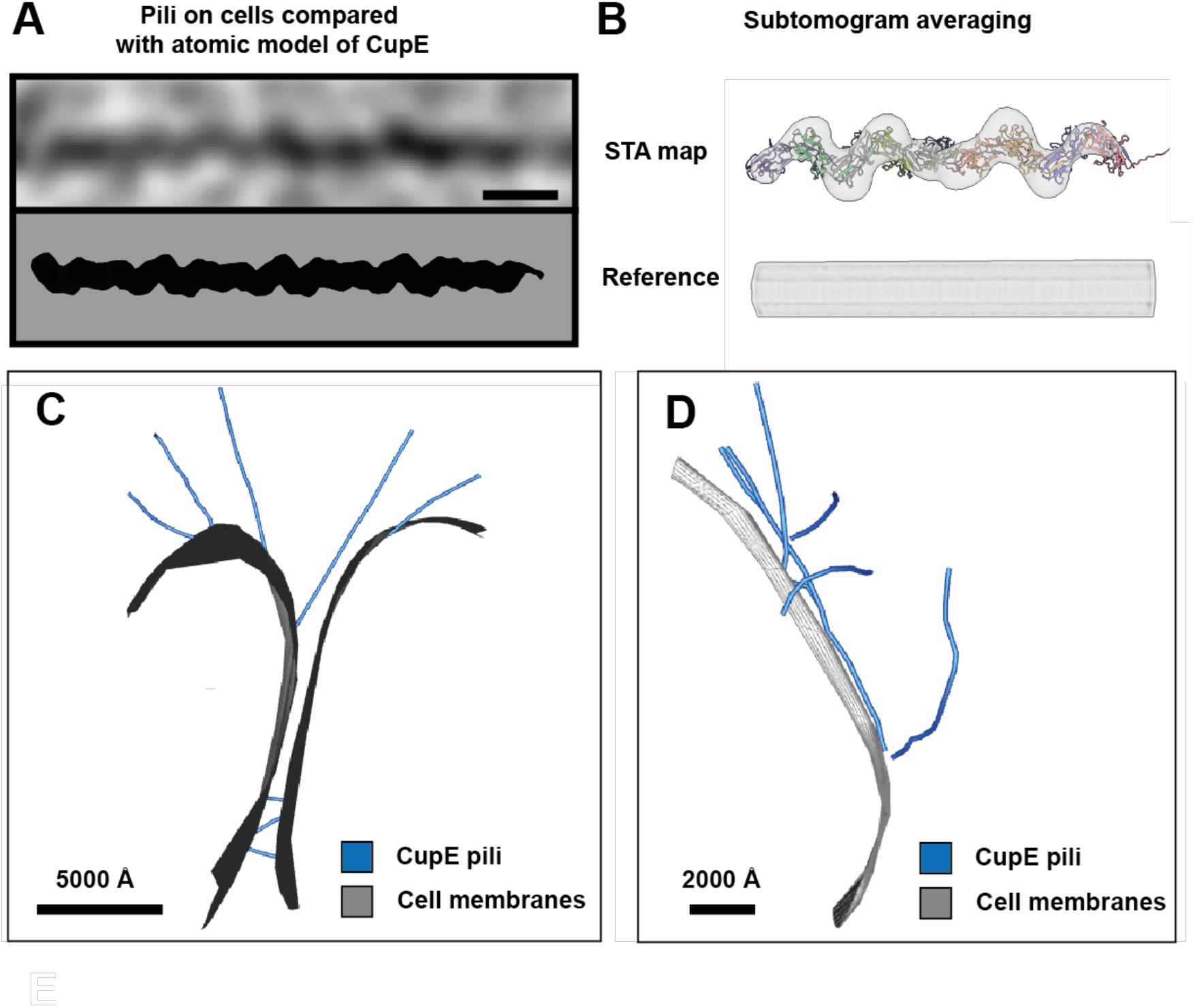
CupE pili imaged on cells recapture features of isolated CupE pili. **(A)** Cryo-ET of pili on cells (upper) recaptures the size and zigzag architecture of the atomic model of CupE, which was projected at 10 Å resolution for comparison (lower). Scale bar is 100 Å. **(B)** Subtomogram averaging of pili on cells results in zigzag-shaped density consistent with the atomic model shown as ribbons. Particles were aligned against a cylindrical reference to prevent bias. **(C-D)** Segmentation of tomograms as shown in Figure 3.

**Figure S6:**
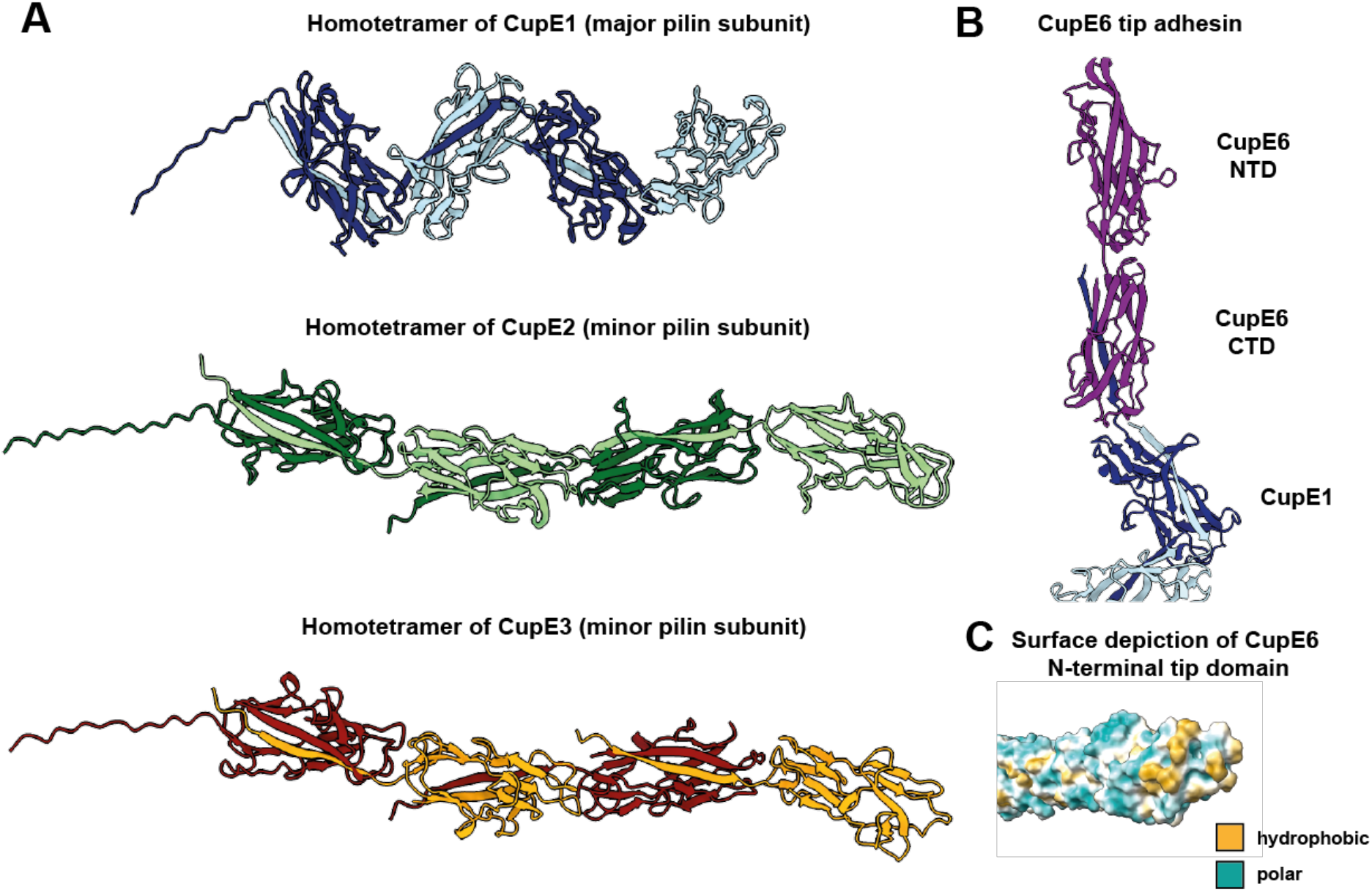
Structural predictions of minor pilins CupE2 and CupE3 and the tip adhesin CupE6 by AlphaFold2. **(A)** Predictions of homotetramers of CupE1, CupE2, and CupE3. The model for CupE1 was validated by comparison with the cryo-EM structure (C_α_-RMSD of E1/E1 subunits: 0.96 Å) (Figures 1 and 2). All pilins of CupE are predicted to share a similar structure (C_α_-RMSD E1/E2: 1.64 Å, E1/E3: 1.63 Å, E2/E3: 0.89 Å), and CupE2 and CupE3 are also predicted to polymerize through donor strand complementation. Pilins mainly differ in domain orientation within the filament. **(B)** Prediction of a filament consisting of three subunits of CupE1 capped with the CupE6 adhesin tip subunit. The adhesin protein CupE6 consists of two domains, with the N-terminal domain capping the filament and the C-terminal domain exhibiting hydrophobic surface patches as predicted in previous studies (Pakharukova et al., 2018). **(C)** CupE6 adhesin domain prediction as in (B) shown as hydrophobic surface depiction, revealing a hydrophobic patch at the domain tip.

**Figure S7:**
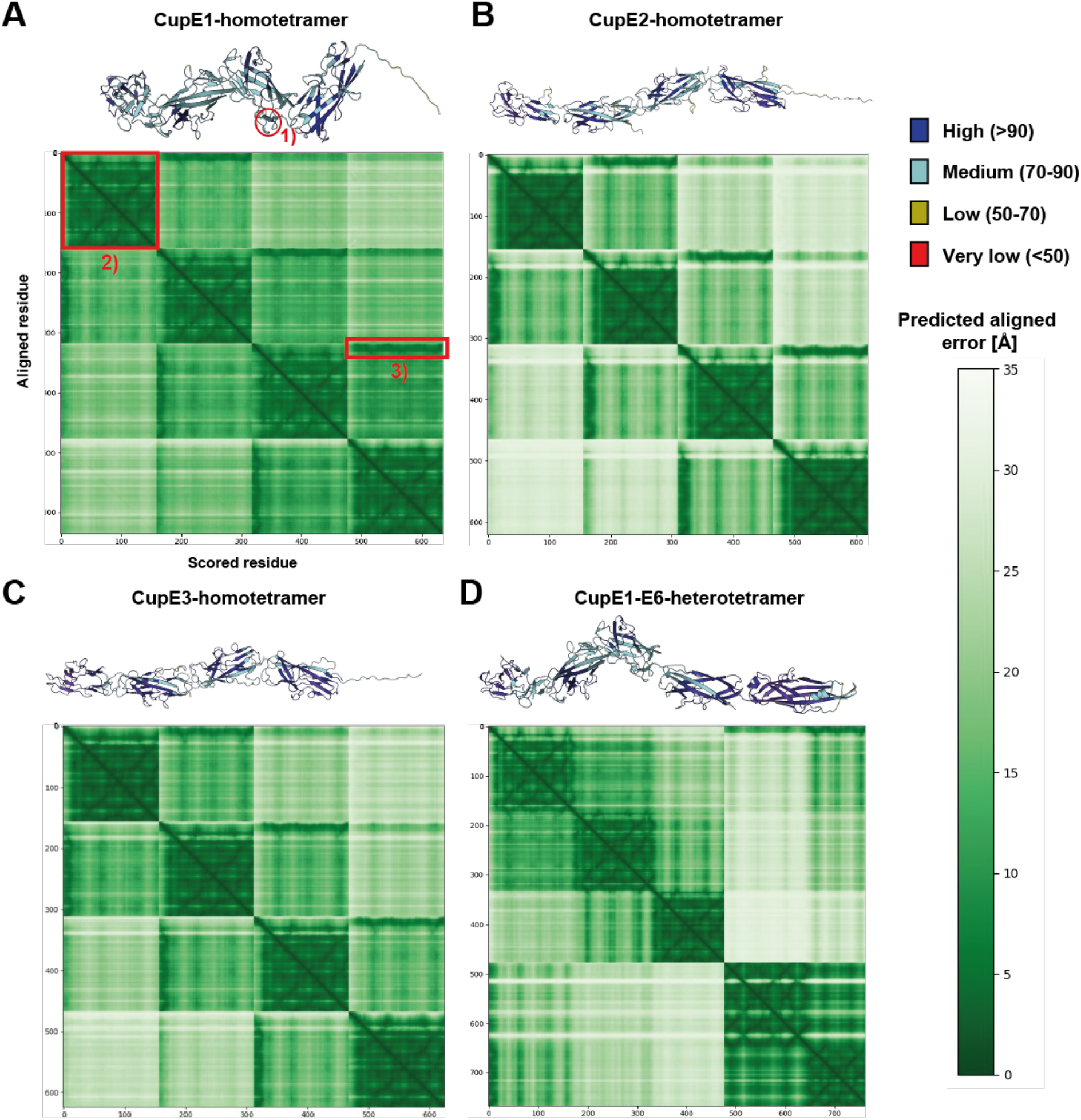
Prediction confidence measures for all tetramers shown in Figure S6. **(A-D)** The prediction confidence is evaluated with different scores. Structures are colored according to their predicted Local Distance Difference Test (pLDDT). The pLDDT measures the model confidence per residue (Jumper et al., 2021). The pLDDT for all tetramers is very high or high for most parts of the prediction and low only for β-hairpin motifs (e.g. labeled as ‘1’ in (A)). The PAE (Predicted Aligned Error) measures the expected positional error (in Å) at residue X, when the predicted and true structures are aligned on residue Y (Tunyasuvunakool et al., 2021, Jumper et al., 2021). The PAE is visualized as a heatmap, where green means low expected error, showing the PAE for every pair of residues, resulting in 4×4 submatrices for every PAE heatmap. Every submatrix (e.g., labeled as ‘2’) on the diagonal shows the intra-subunit PAE. It shows that the intra-domain PAE for all filaments and the PAE for the donor β-strand interaction (e.g., labeled as ‘3’) is low, arguing that the protein structure for one filament subunit is predicted correctly. The inter-subunit predicted error increases when the distance between the subunits is greater. This implies on the one hand that the inter-subunit orientation prediction should not be taken as a precise measurement. On the other hand, it could mean that the filament is flexible and therefore no rigid inter-subunit conformation exists, consistent with our cryo-EM data. In (D), the relative domain position within the CupE6 adhesin is predicted with high confidence, but the PAE for the relative orientation of CupE6 against the CupE1 filament is high. This could again mean high flexibility, but also that the predicted adhesin orientation must be interpreted with care.

## Supplementary Tables

**Table S1:** CupE1 cryo-EM data acquisition and processing statistics.

| **Data collection and processing** | CupE1 WT [EMDB XXXX, PDB XXXX] | CupE1 111-113<sub>AGA</sub> [EMDB XXXX, PDB XXXX] |
| --- | --- | --- |
| Microscope | Krios Titan G3 | Krios Titan G3 |
| Magnification | 81,000 | 81,000 |
| Voltage (kV) | 300 | 300 |
| Electron exposure (e<sup>-</sup>/Å<sup>2</sup>) | 46 | 45 |
| Defocus range (μm) | -1 to -2.5 | -1 to -2.5 |
| Pixel size (Å) | 0.546 (super-resolution)<br>1.092 (physical, final) | 0.546 (super-resolution)<br>1.092 (physical, final) |
| Symmetry imposed | Helical, final: 214.56° 33.12 Å | Helical, final: 214.67°, 33.04 Å |
| Initial particle images (no.) | 3,766,858 | 1,259,369 |
| Final particle images (no.) | 274,457 | 88,886 |
| Map resolution (Å) | 3.47 | 4.13 |
| FSC threshold | 0.143 | 0.143 |
| Map resolution range (Å) | 3.4-4.2 | 3.8-4.7 |
| **Refinement** |  | No atomic model built |
| Initial model used (PDB code) | n/a |  |
| Model resolution (Å) | 2.9/3.1/3.9 |  |
| FSC threshold | 0/0.143/0.5 |  |
| Model resolution range (Å) | n/a |  |
| Map sharpening *B* factor (Å<sup>2</sup>) | -97 | -152 |
| Model composition |  |  |
| Non-hydrogen atoms | 5655 |  |
| Protein residues | 795 |  |
| Ligands | 0 |  |
| *B* factors (Å<sup>2</sup>) |  |  |
| Protein | 66.88 |  |
| Ligand | n/a |  |
| R.m.s. deviations |  |  |
| Bond lengths (Å) | 0.002 |  |
| Bond angles (°) | 0.547 |  |
| Validation |  |  |
| MolProbity score | 1.95 |  |
| Clashscore | 8.03 |  |
| Poor rotamers (%) | 0 |  |
| Ramachandran plot |  |  |
| Favored (%) | 91.08 |  |
| Allowed (%) | 8.92 |  |
| Disallowed (%) | 0 |  |

**Table S2:**
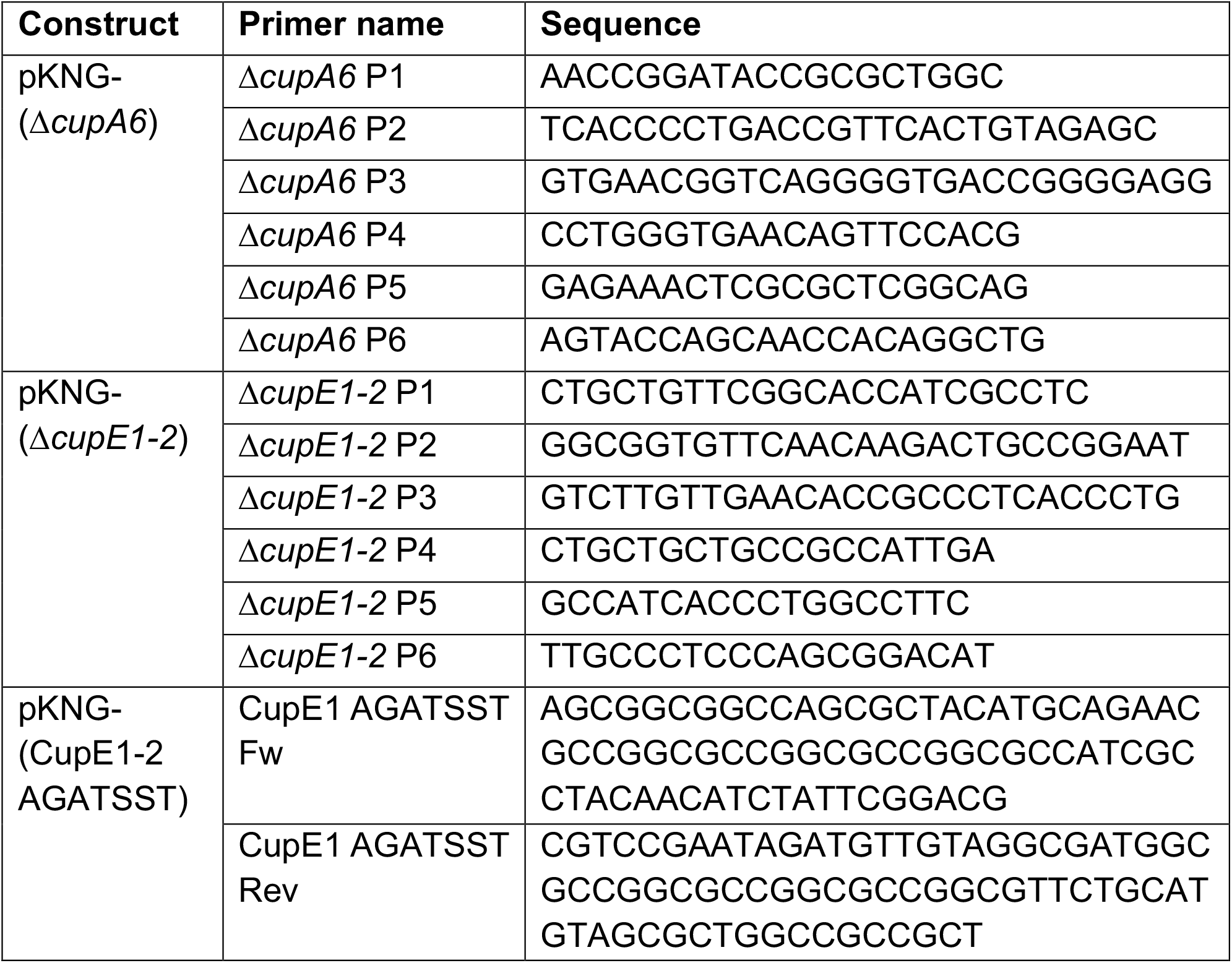
Primers used in this study.

**Table S3:** Plasmids used in this study.

| Plasmid | Characteristics | Source |
| --- | --- | --- |
| Cloning vectors |  |  |
| pCR<sup>TM</sup>-Blunt II-TOPO<sup>TM</sup> | Sub-cloning vector for constructs synthesized by KOD PCR, (Km<sup>R</sup>) | Invitrogen |
| pRK2013 | Self-transmissible helper plasmid for three-partner conjugations, Km<sup>R</sup> | Figurski and Helinski (1979) |
| Chromosomal mutagenesis vectors |  |  |
| pKNG101 | Non-replicative suicide vector for *P. aeruginosa* chromosome mutagenesis, ori6K, mobRK2, *sacB* gene for sucrose sensitivity, Sm<sup>R</sup> | Kaniga et al. (1991) |
| pKNG-(Δ*cupA6*) | Suicide vector to delete *cupA6* (PA2133) from *P. aeruginosa*, Sm<sup>R</sup> | This study |
| pKNG-(Δ*cupE1-2*) | Suicide vector to delete *cupE1-2* (PA4648-9) from *P. aeruginosa*, Sm<sup>R</sup> | This study |
| pKNG-(CupE1-2 AGATSST) | Introduction of mutation of threonine-rich loop TTTTSST to AGATSST in CupE1 | This study |

**Table S4:**
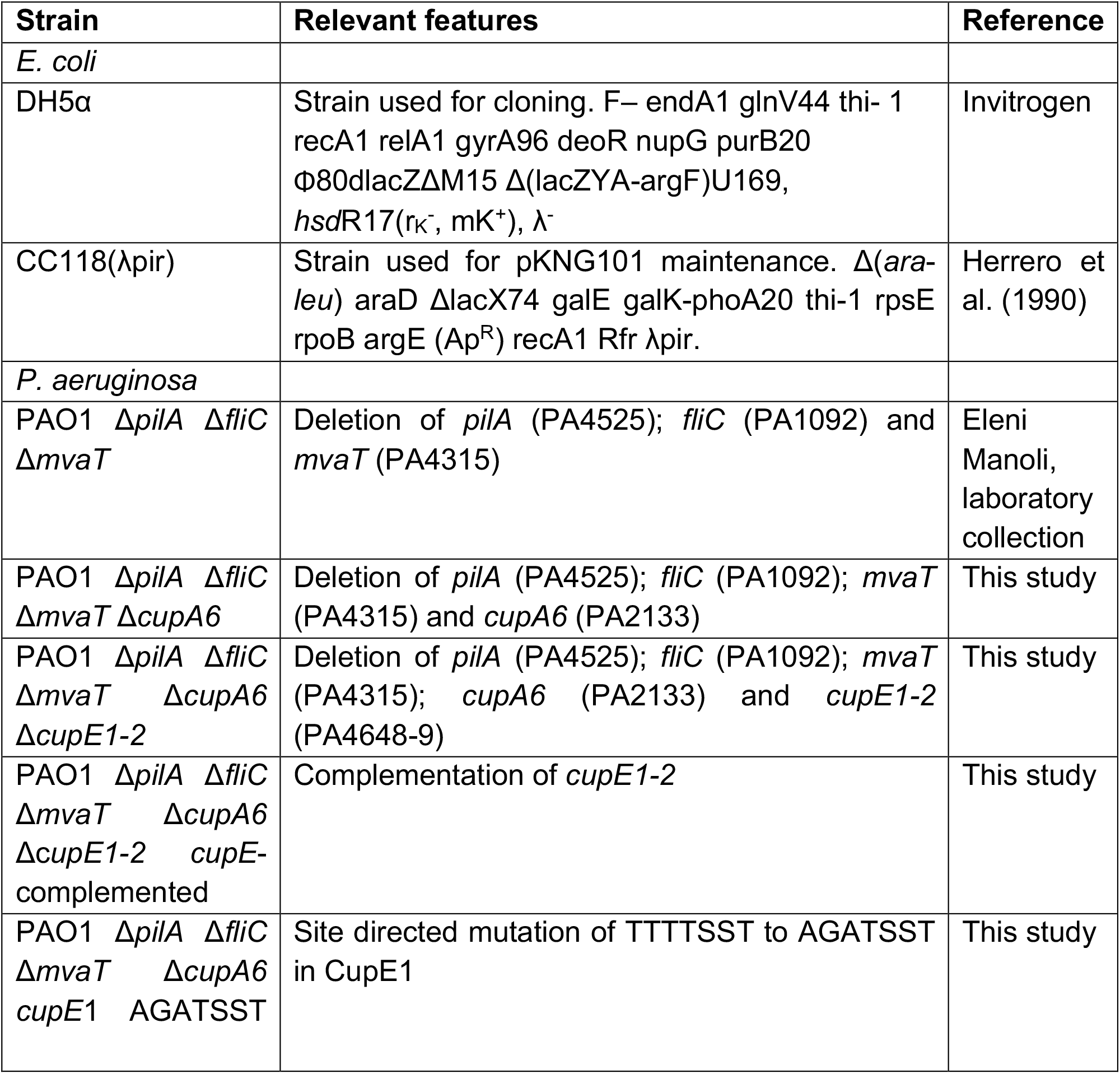
Strains used in this study.

**Table S5:**
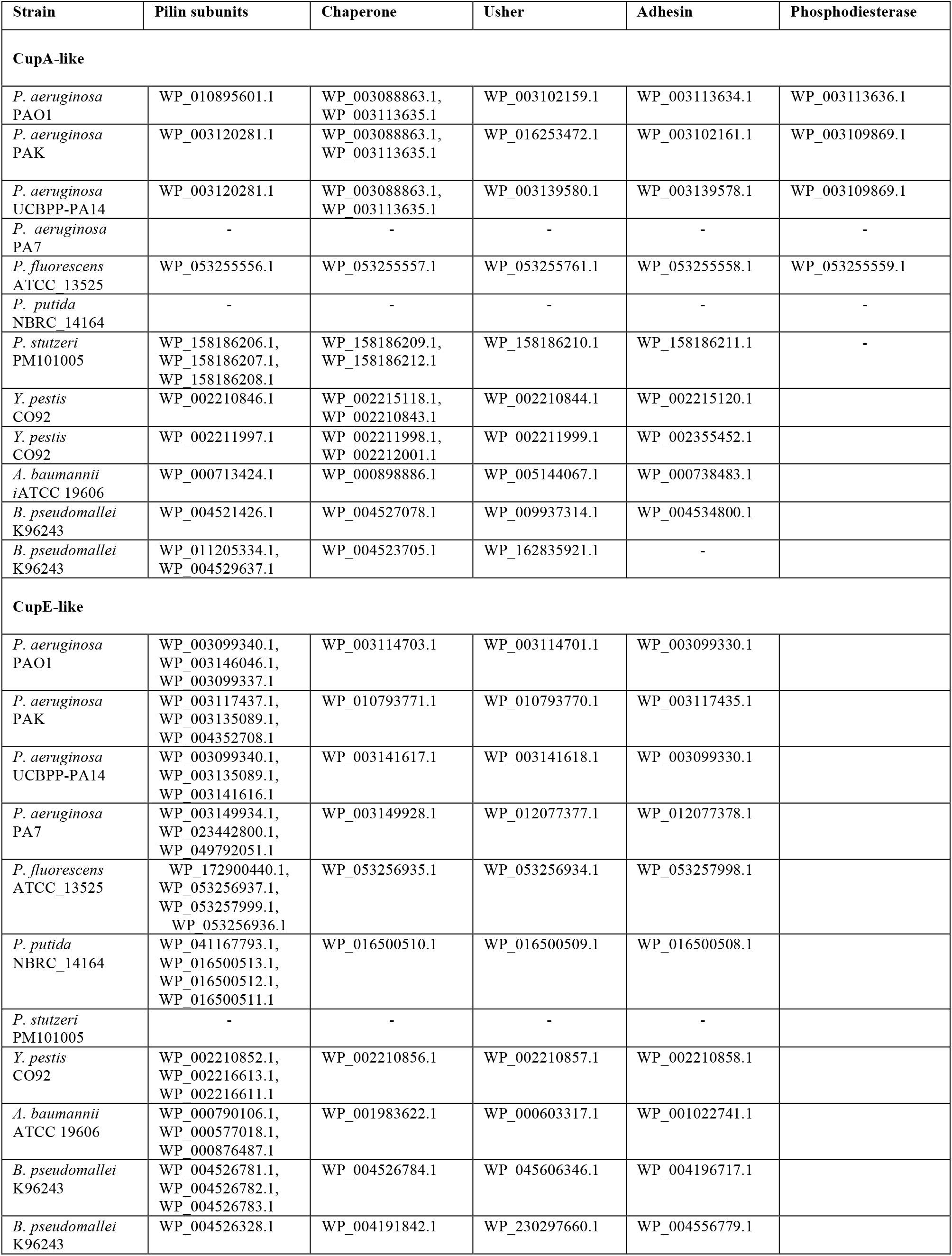
Accession details for proteins encoded by genes shown in Figure 4 and for proteins shown in Figure S4. For the complete assignment of NCBI gene loci to Cup gene clusters for every strain of *P. aeruginosa* of the *Pseudomonas* Genome Database, please refer to the .xlsx file additionally included with the manuscript. A “*” before the location in the supplementary file indicates that only a pseudogene was found. This is likely due to sequencing errors.

## Movie Legends

**Movie S1: Cryo-EM reconstruction of CupE1 from *P. aeruginosa***.

A 3.5 Å resolution cryo-EM density map of the CupE pilus is shown at an isosurface threshold of 15 σ away from the mean, which was used to build an atomic model of the main pilus-forming subunit CupE1. The zigzag-arranged Ig-like domains of CupE1 are complemented and stabilized by donor strand exchange between the individual subunits (surface depiction and ribbon diagrams shown).

**Movie S2: Electron cryotomographic imaging of CupE pili on cells**. Tomographic ‘Z’-slices through *P. aeruginosa* cells and their surrounding region are sequentially shown and reversed when the end of the pilus is reached. CupE pili emerge from the outer membrane of *P. aeruginosa* as extended flexible filaments, here appearing to attach to a particle within their surroundings.

